# The Impact of a Western Diet High in Phosphate on the CKD-MBD in an Alport Syndrome Model

**DOI:** 10.1101/2025.01.17.633378

**Authors:** Matthew J. Williams, Hiral M. Patel, Carley B. Halling, Keith A. Hruska

## Abstract

**Background:** Chronic kidney disease – mineral bone disorder (CKD-MBD) is a syndrome that begins early in CKD, contributes to CKD-associated mortality, and includes components of FGF23 elevation, αklotho deficiency, CKD-stimulated vascular disease, and renal osteodystrophy. Hyperphosphatemia, occurring in later stages of CKD, is also driven by mechanisms of CKD-MBD, and has been shown to stimulate vascular calcification. In a mouse model of Alport CKD that is resistant to vascular calcification, we examine the effects of a high-phosphate Western-type diet on the CKD-MBD, and test whether the diet promotes induction of vascular calcification.

**Methods:** An X-linked Col4a5 deficient murine homolog of Alport Syndrome (CKD) and wild type (WT) littermates were fed an animal protein 1.2% high phosphate diet or a standard vegetable protein diet. At disease progression equivalent to CKD stage 4-5, we examined kidney histology for fibrosis, blood for BUN (marker of CKD), and markers of CKD-MBD disease progression, kidney tissue for klotho production, and aorta histology and tissue mRNA and protein analysis for vascular calcification.

**Results:** The Western high Pi diet produced hyperphosphatemia in the CKD animals compared to WT and increased plasma PTH (1880 from 110 pg / ml), FGF23 c-term (670 from 120 pg / ml), and FGF23 intact (3780 from 280 pg / ml), and reduced kidney klotho mRNA and protein (57-67% reduction) (all p < 0.01). Referenced against the CKD animals fed vegetable-based diet, the Western high phosphate-fed CKD animals showed higher levels of plasma PTH and FGF23s. In the wild-type control mice with normal renal function, Western diet produced increased PTH, intact FGF23, and reduced renal klotho (all p <0.01). Vascular smooth muscle transdifferentiation and vascular calcification was not induced by Western high phosphate diet in this model of CKD.

**Conclusions:** Our results show that a Western-style high-phosphate diet advances elements of the CKD-MBD. Renal klotho, FGF23 and PTH are affected by diet even with normal kidney function, suggesting a need for early intervention in the management of phosphate homeostasis as a component of CKD therapy. Additionally, CKD, klotho, and FGF23 all are associated with early aging. Therefore, our findings suggest that a Western high Pi diet accelerates aging and would contribute to the systemic complications of CKD – cardiac disease, osteodystrophy, and vascular disease.

## Introduction

Chronic kidney disease (CKD) is a public health pandemic affecting ∼850 million people including 5.5 million Americans (Center for disease Control, 2023).(1) Advanced CKD is associated with several adverse clinical outcomes, such as accelerated cardiovascular diseases, kidney failure requiring kidney replacement therapy, mineral bone disease, anemia, metabolic and endocrine abnormalities, and poor quality of life. (2, 3) Patients with CKD stage 3 and 4 have a two- to threefold increased risk of cardiovascular (CV) mortality, and this risk is further increased in patients with CKD stage 5 on dialysis (4, 5). The CV mortality associated with CKD is remarkable in that the excess risk compared to the general population is not due to established cardiovascular disease risk factors (6). Rather, studies over the past two decades have identified non-traditional risk factors as contributing to CKD-associated mortality. These non-traditional risk factors include hyperphosphatemia (7), fibroblast growth factor 23 (FGF23) (8, 9), αklotho deficiency (10–13), CKD-stimulated vascular disease (14–16), and renal osteodystrophy (17). These risk factors are components of the syndrome named the CKD-MBD (chronic kidney disease-mineral bone disorder) (18, 19) in recognition of its role in cardiovascular complications of CKD (5, 20–22). Shortly after the CKD-MBD designation, it was found to be a uniform result of kidney diseases and to begin early in the course of disease (stage 2 CKD) (23–26). At these early stages, the CKD-MBD consists of vascular disease (especially arterial stiffness), osteodystrophy, elevated FGF23, and decreased αklotho (12, 23, 26, 27). As kidney disease progresses, the familiar clinical elements of the CKD-MBD become manifest, and calcitriol deficiency, hyperparathyroidism, hyperphosphatemia, hypocalcemia, and cardiac hypertrophy become prevalent (28–30).

Hyperphosphatemia, a component of the CKD-MBD, is associated with all cause and cardiovascular mortality in CKD (7, 31, 32), and in the general population (33). As early as 1982, control of phosphate has been considered as a potential therapeutic option in CKD progression (34). Phosphate has been shown to stimulate vascular smooth muscle cell (VSMC) calcification (35–38), as a mechanism of secondary cardiac disease. The mechanism of phosphate action in vascular calcification and CKD progression is through signal transduction stimulated by PIT1, a Na/Pi transporter on VSMC and renal tubular cell membranes (39, 40). Pi/PIT1 signaling in VSMC stimulates transdifferentiation to an osteogenic phenotype through activation of the Runx2 gene, the master osteoblast transcription factor (41–43).

We have developed a murine model of Col4A5 deficiency Alport syndrome on the C57Bl6J background that does not express conduit artery disease at CKD equivalency to human stage 4-5 CKD (44). Here, we analyze the effects of a Western diet high in phosphate on the CKD-MBD in Alport syndrome. Secondly, we question whether hyperphosphatemia is sufficient to induce vascular calcification or vascular smooth muscle cell (VSMC) transdifferentiation to an osteoblastic phenotype.

## Methods

### Alport mice

We used the murine homolog of X-linked Alport syndrome, which is a deficiency in the gene for the α5 chain of type IV collagen *Col4a5*, as a model of human CKD (45). Normal males were bred with *Col4a5*^+/−^ females to generate males with (CKD) and without (wild type littermates, WT) *Col4a5* knockout. *Col4a5*^+/−^ female selection was based on production of male progeny without vascular calcification. Hemizygote males developed spontaneous kidney disease and were used to study the CKD-MBD. Mice received either a Western (casein-based) 1.2% high Pi diet (Dyets, Inc., 113321), or a standard (vegetable protein-based) chow diet (LabDiet 5053 - PicoLab^®^ Rodent diet 20, 0.61% total Pi with 0.33% non-phytate-bound) beginning at 75 days old (do) until euthanasia. Euthanasia date for standard diet-fed mice was 225 days of life, but mortality was high. Expecting increased mortality with mice fed Western high-phosphate diet, the planned euthanasia date for this group was shortened to 190 days to conserve animals.

### Tissue collection

Euthanasia of mice was performed under injection anesthesia (xylazine 13 mg/kg BW and ketamine 87 mg/kg BW) or inhalant anesthesia (2% isoflurane). Blood was taken by intracardiac stab, and the kidneys, hearts, and aortas dissected en bloc. Blood samples were processed with BD Microtainer gold SST tubes for serum and lavender K2EDTA tubes for plasma. All blood samples were placed on ice at collection and processed samples were stored at −80 °C or below until use.

### Serum and plasma assays

Serum was analyzed for blood urea nitrogen (BUN), serum calcium, and serum phosphate by Midwest Vet Labs (St. Louis, MO) using a Beckman Coulter AU480 Chemistry Analyzer. Plasma protein levels of intact PTH (60-2305, Quidel Corporation), C-terminal FGF23 (60-6300, Quidel Corporation), and intact FGF23 (60-6800, Quidel Corporation) were determined by ELISA.

### Kidney fibrosis and cellularity

Picrosirius red staining was used to determine progression of interstitial fibrosis. Kidney tissues were fixed in 10% neutral buffered formalin overnight, then transferred to 70% ethanol at 4 °C and embedded in paraffin. Embedded tissues were sectioned on the midline frontal plane to 5 micron thickness and stained using the Picrosirius red method. Slides were deparaffinized in xylene and rehydrated in a graded ethanol series. Nuclei were stained with Wiegert’s hematoxylin for 8 minutes, followed by slide wash in tap water for 10 minutes. Slides were then stained with 0.1% (w/v) Direct Red 80 (Millipore Sigma, 365548) in saturated aqueous picric acid (Millipore Sigma, P6744) for one hour followed with 2 washes in 0.5% (v/v) aqueous acetic acid. Finally, the slides were dehydrated with three 1 min washes in 100% ethanol before clearing in xylene and mounting. All histology processing and staining was performed by the Musculoskeletal Research Center Histology Core at Washington University in St. Louis.

Quantitation of fibrosis was determined using fraction of red-stained area in total renal parenchyma. Whole tissue sections were imaged under bright field illumination at 20x in same session at the Hope Center Alafi Neuroimaging Lab using a Hamamatsu NanoZoomer 2.0-HT System with NDP.scan 2.5 software. Images were scaled, converted to 24-bit RGB, and processed using ImageJ FIJI distribution (46). Regions of interest were then scribed to include total area of the tissue. The processing technician was blinded to randomized samples. Area fraction of positive staining was calculated using color thresholding plug-in and the hue, saturation, and brightness (HSB) color model. Parameters were set to isolate red-stained area that increases in fibrosis. Quantitation of cellularity was determined using a similar method. In this case, color determination was set to capture dark-blue-stained cell nuclei.

### RNA preparation and real-time PCR

Total RNA from tissue was isolated and purified using TRIzol (15596018, ThermoFisher Scientific) with tissue disruption by the TissueLyser II (Qiagen), Phasemaker tubes (A33248, ThermoFisher Scientific), and PureLink RNA Mini Kit (12183018a, ThermoFisher Scientific) with PureLink DNase (12185010, ThermoFisher Scientific) according to manufacturers’ instructions. RNA concentration and quality was determined using the NanoDrop™ OneC spectrophotometer (ThermoFisher Scientific). Complementary DNA was generated for real-time PCR (qPCR) using the High-Capacity cDNA Reverse Transcription Kit (4368814, ThermoFisher Scientific). Quantitative PCR analysis was performed with the StepOnePlus Real-Time PCR System (Applied Biosystems) using the PowerUp SYBR Green Master Mix (A25742, Applied Biosystems) run in fast cycling mode. Expression levels were normalized by standardizing cDNA concentration. All biological samples were run using 2 technical replicates. All primers for qPCR were purchased from the LabReady Oligo Service from Integrated DNA Technologies, Inc. (Coralville, Iowa, USA). The list of primers is provided in **Table 1**.

**Table 1.**
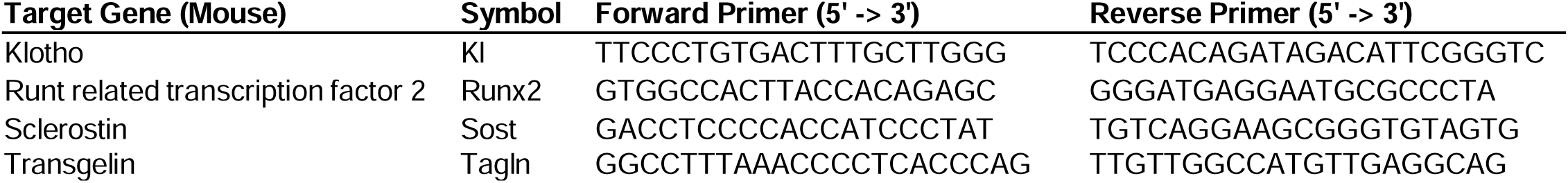
Quantitative PCR primers.

### Aortic calcification

Calcium phosphate in aortas was visualized with Von Kossa histology staining. Freshly excised aortas were cleaned of perivascular adipose tissue, flash frozen in liquid nitrogen, and stored at −80 °C. Once all the frozen aortas in the experimental groups were collected, they were briefly thawed at 4 °C, straightened, immersed in cold (on ice) 10% formalin, fixed overnight at 4 °C, transferred to 70% ethanol, and embedded in paraffin. Embedded aortas, oriented to capture the full length of the tissue, were sectioned twice at 5-micron thickness, at 50-micron difference in depth. Following deparaffinization and hydration, Von Kossa’s stain method progressed with exposure to 5% silver solution (Millipore Sigma, 209139) under bright light for 1 hour, 4 rinses with distilled water, 5-minute exposure to 5% sodium thiosulphate, tap water wash with distilled water rinse, counterstain with Nuclear Fast Red (Millipore Sigma, N3020) for 5 mins, wash in tap water, slide dehydration in a graded ethanol series, clear, and coverslip. All histology processing and staining was performed by the Musculoskeletal Research Center Histology Core at Washington University in St. Louis. Whole aorta sections were imaged under bright field illumination at 20x in same session at the Hope Center Alafi Neuroimaging Lab using a Hamamatsu NanoZoomer 2.0-HT System with NDP.scan 2.5 software. Scanned slides were examined by technician using NDP.view 2.7.52, blinded and randomized.

### Protein extraction and immunoblot analysis

Full length frozen aortas were homogenized in ice-cold RIPA Lysis and Extraction Buffer (ThermoFisher Scientific, 89900) with Protease Inhibitor Cocktail (Millipore Sigma, P8340) and PhosSTOP phosphatase inhibitor (Millipore Sigma, 4906845001) using a TissueLyser II (Qiagen). Total protein of each sample lysate was determined with the Pierce™ BCA Protein Assay Kit (ThermoFisher Scientific). Lysates were then prepared under reducing conditions and loaded with equivalent total protein into a 10% TGX gel (Bio-Rad) for protein separation using gel electrophoresis. Proteins in the finished gel were transferred to a 0.45 PVDF membrane using a Trans-Blot Turbo (Bio-Rad) according to manufacturer’s guidelines. Resulting blots were blocked with 1% milk (Bio-Rad, 1706404) or Li-Cor Intercept Blocking Buffer (927–600001) in TBS and probed with antibodies to 1:1000 Runx2 (D1L7F) (Cell Signaling Technology Cat# 12556, RRID:AB_2732805) or 1:500 Klotho (KM2076) (Cosmo Bio Cat# KAL-KO603, RRID:AB_3390355) in addition to housekeeping control 1:500 Beta-actin (C4) Alexa Fluor 680 (Santa Cruz Biotechnology Cat# sc-47778 AF680, RRID:AB_626632). IRDye 800CW donkey anti-rabbit (LI-COR Biosciences Cat# 926-32213, RRID:AB_621848) and IRDye 800CW goat anti-rat (LI-COR Biosciences Cat# 926-32219, RRID:AB_1850025) at 1:5000 were used as secondary antibodies. Near-infrared fluorescence imaging of multiplexed blots was conducted by the Bio-Rad ChemiDoc MP and quantitative analysis was performed using Bio-Rad Image Lab 6.1.

### Statistics

Four experimental groups are evaluated in this study: wild type (WT) and Alport CKD mice fed a high Pi Western-type diet, and WT and Alport CKD mice fed a standard vegetable-based mouse diet. All data were examined with Shapiro-Wilk test for normality and Levene’s test for equality of variance across groups. Significant differences between two groups with normal distribution and equal variance between groups was determined with Student’s t-test. Significance differences between data sets with more than two groups, approximate normal distributions, and equal variance across groups were determined by ANOVA with Tukey’s multiple comparison test. Significance differences between data sets with more than two groups, approximate normal distributions, and unequal variance across groups were determined by Welch’s ANOVA with Dunnett’s T3 multiple comparison test. Otherwise, significance was determined using Kruskal-Wallis H test with Bonferroni correction for multiple pairwise comparisons. Rare extreme outliers as identified using the interquartile range (IQR) method in SPSS were removed from calculations for significant differences. Comparisons were made between WT and CKD groups to show effect of chronic kidney disease, and between Western diet and vegetable diet groups to show effect of diet change. Analyses of plasma protein levels (**Figure 3C-E**) were performed on log10-transformed data. All quantitative data are presented as mean ± SD and p < 0.05 was considered statistically significant. All data analyses were carried out using IBM SPSS Statistics 29.

## Results

Alport CKD mice in the standard vegetable diet group experienced higher than expected mortality (43%) before planned euthanasia date of 225 days old (do) (**Table 2**). We expected a further increase of mortality with mice fed Western animal protein-based 1.2% high phosphate diet. Therefore, to conserve animals, we reduced euthanasia date for this group to 190 do. At this endpoint, the CKD mice fed a Western-type diet had a 5% mortality (**Table 2**). **Figure 1** shows serum BUN levels are significantly increased in Alport CKD compared to WT littermates for 190 do mice fed the Western high Pi diet and the 225 do mice fed the standard vegetable-based diet. Kidney pathology showed a severe fibrosis and cellular infiltration from the tubulointerstitial nephritis in the 190-day old Western high Pi diet-fed CKD animals (**Figure 2A-B**). Tubulointerstitial nephritis ending in fibrosis is the pathologic hallmark of Alport syndrome kidney disease. (47) Fibrosis and cellular infiltration was also confirmed for the 225 do CKD mice fed the vegetable-based diet (**Figure 2C-D**). Notably, cellular infiltration was significantly increased in the WT animals fed the Western high Pi diet in comparison with those fed the vegetable-based diet. **Figure 2E** shows that the measurement of relative fibrosis is strongly correlated with BUN measurements in Alport CKD mice. Alport syndrome, and as represented in this model, is variable in disease progression. At 190 do, some COL4A5 deficient animals did not show significant elevations of BUN to represent advanced disease (**Figure 1**), and had little fibrosis. In addition, a few wild type animals showed an increase of BUN suggesting early disease initiation. To assure development of CKD and study the contribution of Western type high-phosphate diet on the CKD-MBD, the data in the remaining figures are selected from animals with BUN > 43 for Alport CKD, or BUN < 35 for WT littermates.

**Table 2.** Mortality of mice fed standard chow or the Western high-phosphate diet.

| **Diet** | **Group** | **Days of Life** | **Mortality** |
| --- | --- | --- | --- |
| Standard chow | WT | 225 | 0% (0/25) |
| Standard chow | CKD | 225 | 43% (12/28) |
| Western high-phosphate | WT | 190 | 2% (2/116) |
| Western high-phosphate | CKD | 190 | 5% (8/159) |

Mortality includes mice that have died before targeted euthanasia due to disease or were euthanized before specified euthanasia date due to severe disease. Mice that died due to non-disease causes were not included in total. CKD: Alport col4a5<sup>-</sup>, standard chow diet: vegetable-based protein (0.6% phosphate); Western high-phosphate diet: animal-based protein (1.2% phosphate), WT: wild type littermates to CKD mice.

**Figure 1.**
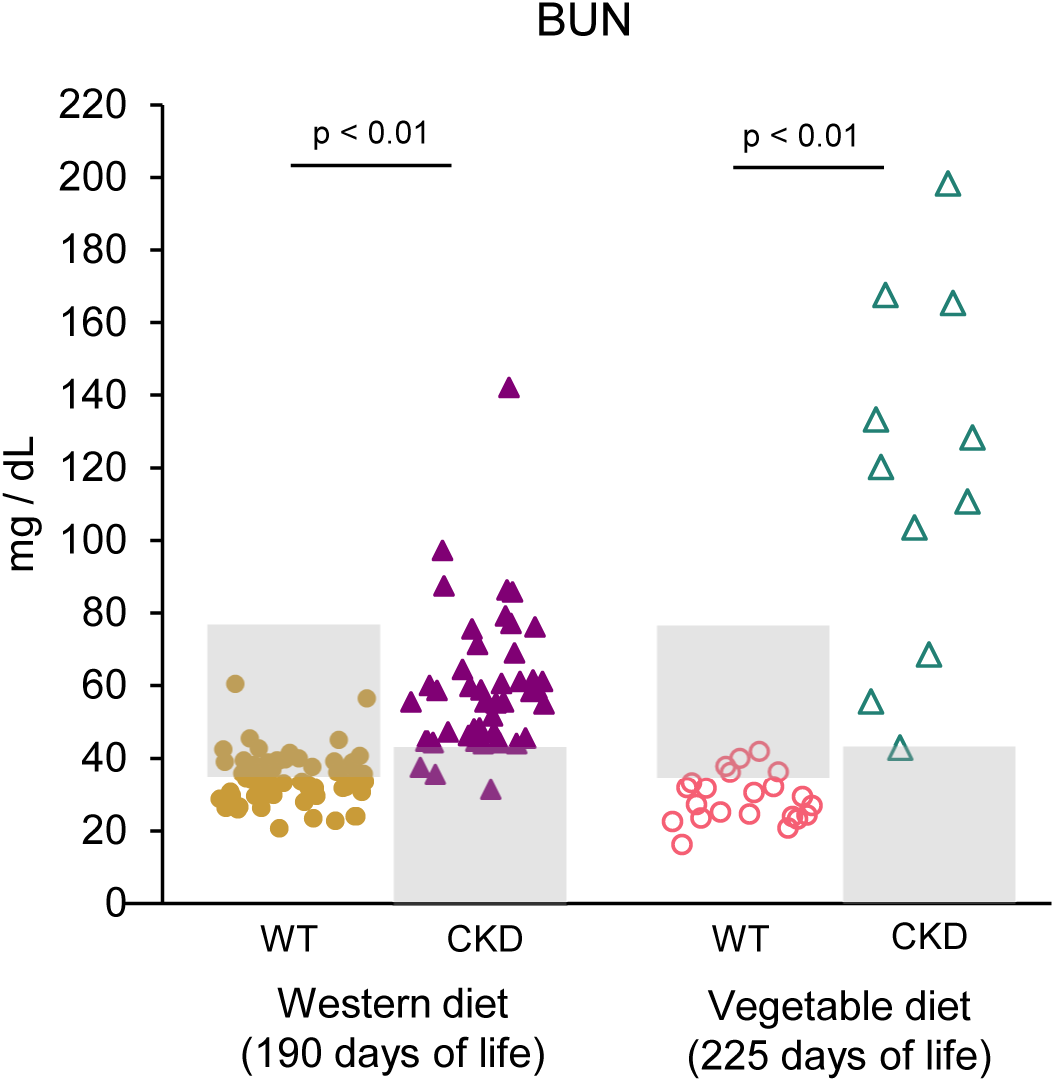
Blood urea nitrogen (BUN) levels in Alport CKD mice fed Western type, high-phosphate (Pi) animal-based protein diet or standard, vegetable-based protein diet compared to their wild-type (WT) littermates. Western diet: WT, n = 62; CKD, n = 45. Vegetable diet WT, n = 22; CKD, n = 11. Shaded areas indicate study animals not selected for normal kidney function in WT (BUN > 35) or confirmation of disease in CKD (BUN < 43). Data are represented as means ± SD. Significance differences between groups was determined using Kruskal-Wallis H test with Bonferroni correction for multiple pairwise comparisons.

**Figure 2.**
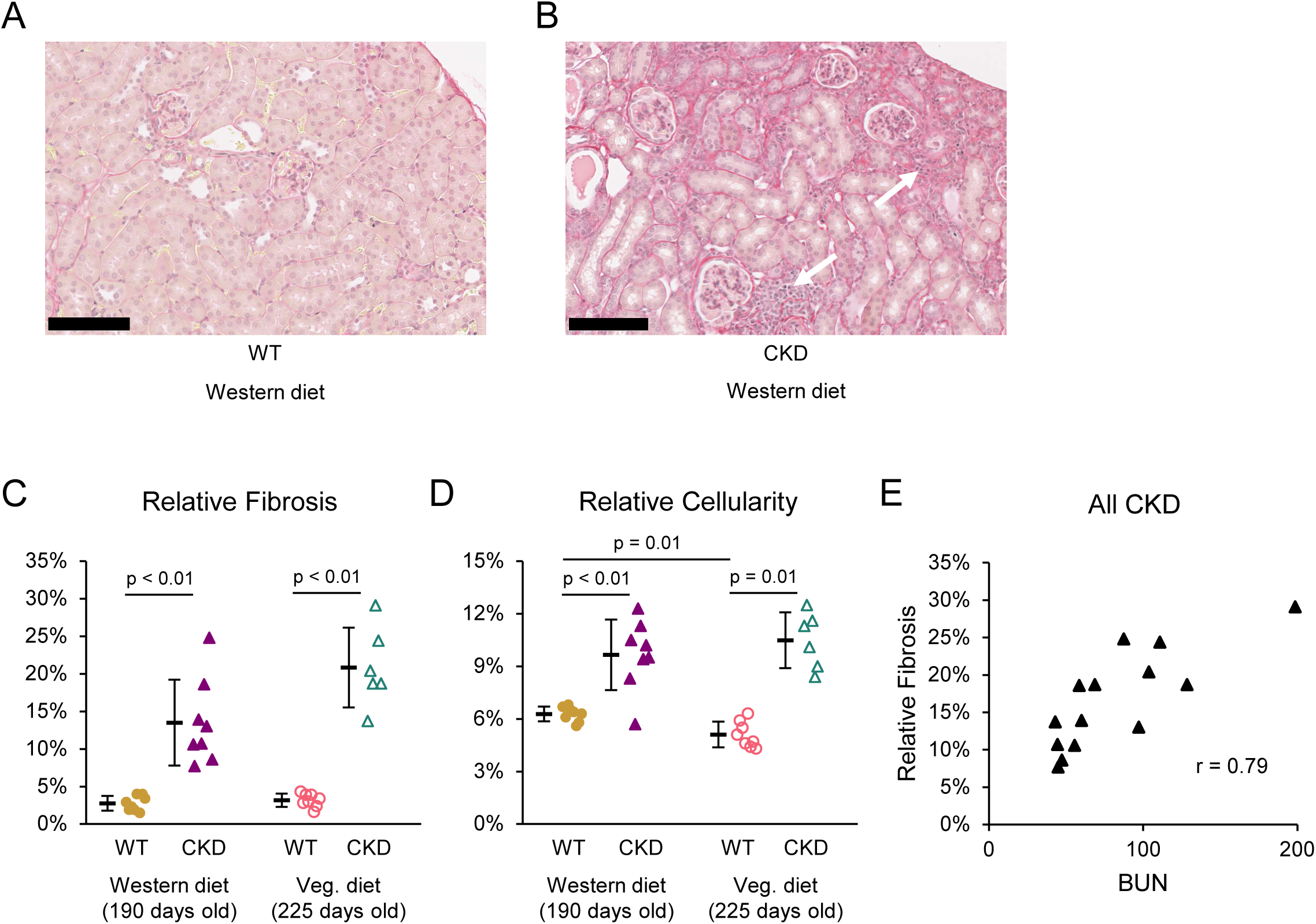
Renal fibrosis in the Alport mice. **A**: Representative picrosirius red-stained kidney section from WT littermates fed the Western-type high Pi diet. **B**: Representative kidney section from 190 day old (do) Alport CKD mice fed the Western-type high Pi diet. Areas of increased interstitial cellularity are indicated with white arrows. **C**: Quantitation of renal fibrosis calculated by total area fraction of red stain in whole tissue images. The four groups of mice include 190 do littermates (WT) fed Western-type high Pi diet; Alport CKD mice fed Western-type high Pi diet, 225 do WT littermates fed standard vegetable-based chow; and Alport CKD mice fed standard chow. **D**: Quantitation of interstitial cellularity in CKD by increased area fraction of nuclear stain. **E**: Correlation between BUN and renal fibrosis measurement in all CKD mice. **A-B**: Bar is 250 microns. **C-D**: Group sizes were n = 6-8 for fibrosis and cellularity scoring. Data are represented as means ± SD. Significant differences between groups were determined using Welch’s ANOVA with Dunnett’s T3 multiple comparison test.

The Western-type high Pi diet produced hyperphosphatemia and hypocalcemia in the Alport CKD animals (**Figure 3A-B**). Plasma levels of the CKD-MBD components, PTH and FGF23, are shown in **Figure 3C-E**. PTH levels were significantly increased in the 190 do high Pi-fed Alport CKD mice to 1880 pg / ml from 110 pg / ml in the WT mice (**Figure 3C**). In 225 do normal Pi-fed mice, PTH levels in the Alport CKD mice were 850 pg / ml compared to 30 pg / ml in the WT littermates. The high Pi diet even increased PTH levels in the WT littermates when referenced against the normal Pi fed WT mice. C-terminal FGF 23 levels were elevated in Alport CKD high Pi diet-fed mice (**Figure 3D**) to 670 pg / ml from 120 pg / ml in the WT littermates. C-terminal FGF23 levels in the normal Pi-fed Alport CKD mice were 160 pg / ml compared to 100 pg / ml in the WT littermates (**Figure 3D**). FGF23 intact hormone levels were increased to 3780 pg / ml in Alport CKD mice from 280 pg / ml in WT mice fed Alport CKD high Pi diet (**Figure 3E**). Intact FGF23 hormone levels in the normal Pi fed Alport CKD mice were 640 pg / ml compared to 120 pg / ml in the WT littermates (**Figure 3E**). The Western high Pi diet worsened the FGF23 arena in the CKD mice and even increased FGF23 levels in the WT littermates referenced against the normal Pi fed WT mice.

**Figure 3.**
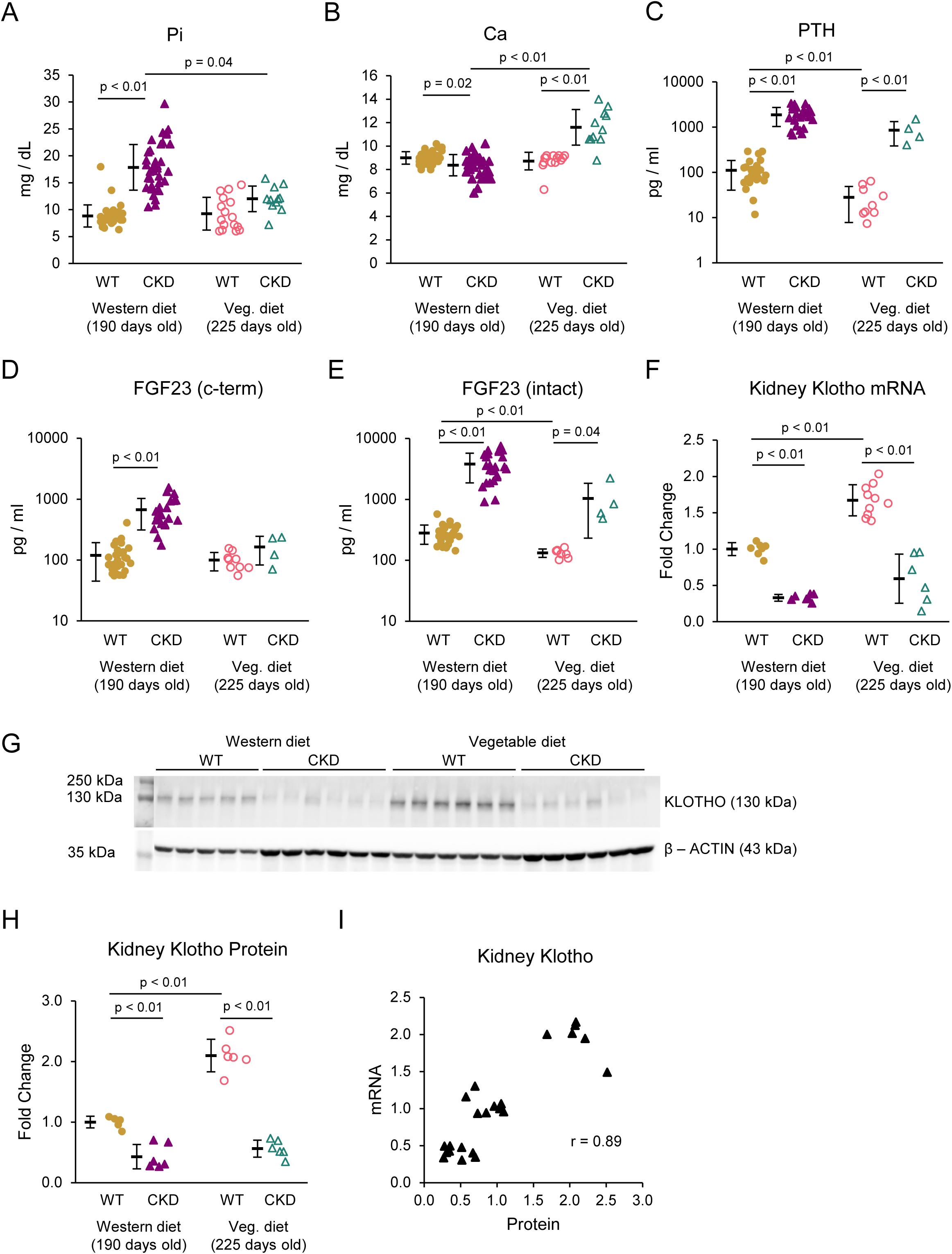
Serum chemistry, plasma proteins, and renal klotho for the chronic kidney disease – mineral bone disorder (CKD-MBD). Groups include WT and Alport CKD mice fed high-phosphate Western diet or vegetable-based diet. **A**: Serum inorganic phosphate (Pi) is significantly increased in CKD with mice on high phosphate Western diet. **B**: Blood calcium (Ca) shows a mild decrease in CKD with mice on a high-phosphate diet and an increase in CKD with mice on standard diet. **A-B**: Western diet groups n = 35-42; vegetable diet groups, n = 11-16. Plasma levels of PTH (**C**), FGF23 (c-term) (**D**), and FGF23 (intact) (**E**) are all significantly elevated in diseased animals on high-phosphate diet compared to their wild type littermates on the same diet. PTH (**C**) and FGF23 (intact) (**E**) also show increase in CKD on normal-diet-fed mice, although at levels reduced compared to counterparts on high-phosphate Western diet. **C**-**E**: Western diet: WT, n = 28; CKD, n = 24. Vegetable diet WT, n = 9; CKD, n = 4. Significance differences between groups was determined using Kruskal-Wallis H test with Bonferroni correction for multiple pairwise comparisons. Data displayed on log10 scale. **F**: Relative kidney klotho mRNA levels of WT and CKD mice fed either high-phosphate Western diet or vegetable-based standard diet (n = 6-10). Klotho mRNA expression normalized to HPRT. Significant differences between groups were determined using Welch’s ANOVA with Dunnett’s T3 multiple comparison test. **G**: Immunoblot of kidney klotho protein (n = 5-6) with relative quantification (**H**). All protein samples were derived from the same kidneys used in klotho mRNA analysis. Significant differences between groups were determined using ANOVA with Tukey’s multiple comparison test. Correlation between mRNA and protein klotho levels is strong (**I**). Data are represented as means ± SD. All data limited by animals with associated BUN <35 for WT and >43 for CKD. CKD, chronic kidney disease; FGF23, fibroblast growth factor 23; PTH, parathyroid hormone; WT, wild type.

Kidney αklotho mRNA levels in kidney homogenates of Alport mice fed the Western high Pi diet were reduced compared to WT (**Figure 3F**). The reduction in αklotho was relatively equivalent to that observed in the 225 do regular chow fed Alport mice (**Figure 3F**). However, comparison of αklotho levels between the WT groups (**Figure 3F**) revealed that the High Pi Western diet reduced renal klotho levels in wild type mice. This effect was confirmed with tissue protein analysis (**Figure 3G-H**), and correlation between mRNA and protein measurements were strong (**Figure 3I**) Soluble circulating klotho (sklotho) levels were not determined, due to the lack of validation and comparability in various murine ELISAs. We did not possess the serum stock to independently validate klotho ELISAs.

### Vascular (aortic) Ca and gene expression in Alport mice

Vascular calcification detected by von Kossa staining of aortic sections from 190 do Alport CKD mice fed the Western high Pi diet was absent (**Figure 4A**). Aortic *Runx2* mRNA levels (the key marker of VSMC transdifferentiation to osteoblastic phenotype) were not increased (**Figure 4C**), nor were aortic *Sost* levels (**Figure 4D**). Furthermore, aortic *Tagln* (sm22α) mRNA levels (a marker of VSMC differentiation) were not decreased (**Figure 4B**). Runx2 protein levels as detected by Western analysis were undetectable in aortas from Western high Pi-fed WT and Alport CKD mice (**Figure 4E**).

**Figure 4.**
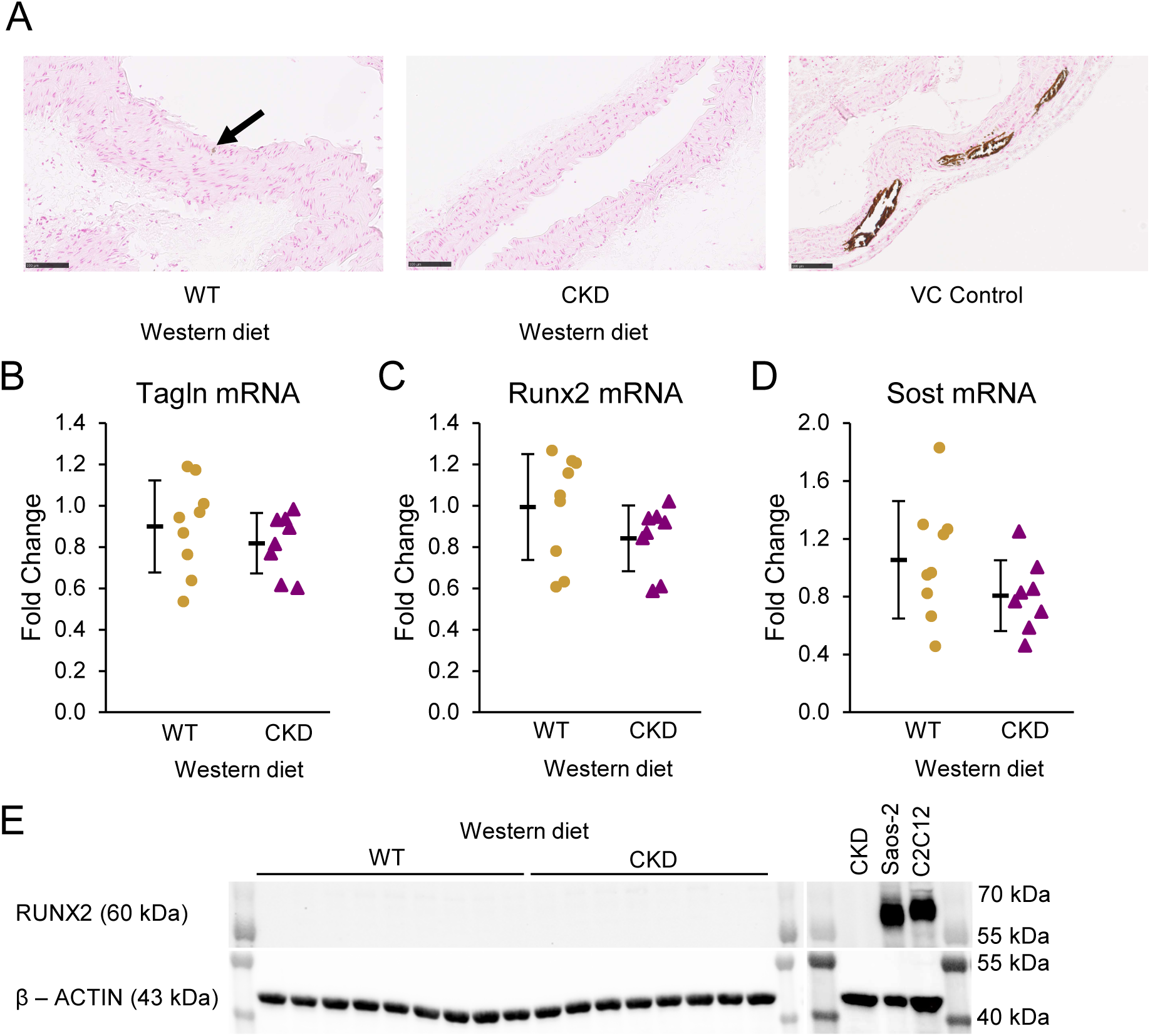
Vascular calcification (VC) in Alport CKD mice fed Western high-phosphate diet. **A**: Histological images of aortas with Von Kossa stain. Both WT and CKD animals showed no significant evidence of vascular calcification (n = 10) as compared to aortic tissue from a mouse model of vascular calcification (DBA/2J mouse with CKD and VC induced by 6 weeks of 0.2% adenine followed by 6 weeks of 0.2% adenine and 0.9% high-phosphate diet). There was presence of approximately 0-3 areas of punctate, <10 micron points of positive stain in full length of axial aorta in both WT and CKD mice (black arrow). Bar is 100 microns. **B-D**: Aortic mRNA from whole tissue lysates of WT and CKD mice fed high-phosphate diet (n = 8-9). *Tagln* (transgelin, or sm22a), a marker of vascular smooth muscle cell differentiation (**B**), *Runx2* (runt-related transcription factor 2), a marker of osteoblastic transition (**C**), and *Sost* (sclerostin), an osteocytic marker (**D**), all show no significant change of overall expression between groups. All target genes are displayed normalized to *rplp13*. Data are represented as means ± SD. **E**: Western blot showing no Runx2 protein expression in whole aorta lysates of WT and CKD mice on high-phosphate diet (n = 8-9). Saos-2 and C2C12 osteoblastic cell lines are shown as positive controls for Runx2. Saos-2 is loaded in equivalent mass to the samples. BETA-ACTIN (multiplexed with RUNX2) was used as a housekeeping protein for normalization. A-E: Includes data limited by animals with associated BUN <35 for WT and >43 for CKD. CKD, chronic kidney disease; WT, wild type.

## Discussion

The data reported here show that changing from a vegetable protein-based diet to an animal protein high phosphorus Western-type diet worsens the severity of the CKD-MBD. On the Western-type high phosphorus diet, Alport CKD mice showed increased blood phosphorus, PTH, and FGF23 levels, decreased blood calcium, and decreased kidney klotho levels. The increases in blood phosphorus, the hypocalcemia, and FGF23 levels in CKD were worse with mice on the Western-type high phosphorus diet at 190 days old than on the vegetable protein-based regular laboratory chow at 225 days old. Even in WT littermate mice with normal kidney function, the Western-type high phosphorus diet negatively affected PTH, intact FGF23, and kidney klotho levels when referenced against the older vegetable protein-fed mice. The effects of the Western-type high phosphorus diet on increasing PTH and FGF23 levels in WT mice probably represent adaptations to maintain phosphate homeostasis and normal blood phosphorus levels. However, the reduction in renal klotho levels is potentially even more important. Reductions in klotho induce resistance to FGF23 signaling.(48) By inducing FGF23 resistance, the mice were “primed” for the effects of CKD, resulting in the increased severity of the CKD-MBD in the Western-type high phosphorus fed CKD mice. Since phosphorus, (49) FGF23, (50) CKD (51, 52) and klotho (53, 54) are all factors contributing to aging, it is possible that they all work in aging through the functions of klotho.(55) Since klotho reductions are among the earliest perturbations in the development of the CKD-MBD, (56) it looms as an attractive target for intervention in the CKD-MBD syndrome.

These results are consistent with prior studies (57, 58). They accentuate the key role of phosphate in our animal-based protein Western diet. This suggests that avoiding processed foods using phosphate-based preservatives (which are similar to the Pi addition in our diet) and substituting vegetable protein for meats when possible, would provide efficacy to the treatment of CKD and the CKD-MBD (59, 60). Further, given that SGLT2 inhibitors worsen CKD-MBD (61–63), including increase of serum phosphate and FGF23, the results herein open the opportunity to test whether diet modification improves efficacy of SGLT2 agents in slowing progression of CKD. We show that the Western-type high phosphate diet affected renal klotho levels independent of kidney function. Since phosphate, FGF23, PTH, CKD, and klotho have been suggested to be premature aging-associated factors, we suggest that klotho deficiency could be the unifying pathophysiologic mechanism tying all of the factors together (64). Our klotho data suggest that the aging-like effects of CKD may be mediated through our diet, that the fracture prevalence of CKD may relate to dietary effects on PTH levels, and that the cardiovascular effects of CKD may relate to dietary effects on FGF23 and klotho (65).

We previously reported the development of an Alport CKD model without vascular calcification, vascular stiffness, or hypertension (44). In the present study, the addition of a Western type high-phosphate diet to this model did not promote smooth muscle transdifferentiation and vascular calcification. This is despite the observed hyperphosphatemia, and the role of phosphate to affect the Na-Pi cotransporter Pit-1 in renal proximal tubules and vascular smooth muscle cells (66). The resistance of the C57Bl6J strain background to vascular calcification was not overcome. This provides an opportunity to study the progression of CKD-MBD pathophysiology without the contribution of vascular calcification. We have used the opportunity to show that kidney disease and the CKD-MBD directly affect cardiac metabolism independent of vascular disease. (44)

There are weaknesses in the data presented here. Measurement of soluble klotho has been fraught with lack of adequate validation of the multiple ELISA assays available. For instance, the most widely used ELISA (IBL International, Japan) was not validated during an analysis of klotho in the CRIC studies. (67) This calls into question the accurate detection of purported klotho proteins by the ELISAs in global klotho knock out mice, and contributes to the lack of comparability between various assays. (68) At the time of this manuscript preparation, we did not have sufficient serum to readdress the issue and independently validate the ELISAs currently used. Secondly, we did not investigate calciprotein particles in our mice with absence of conduit arterial disease to see if these were characteristic of non-calcifying particles. Furthermore, because we had already reported that the mice without conduit arterial disease had a milder renal osteodystrophy (44), not harboring the abnormal osteoblastic activity that is an important component of excessive parathyroid hormone activity and CKD osteodystrophy (69), we deferred on assessing the osteodystrophy of the cohorts reported herein. Thus, the bone-vascular paradox was not interrogated. Despite the robust changes in FGF23 that we observed, we did not assess anemia and iron deficiency as contributing to this aspect of the CKD-MBD (70).

In summary, our data provide strong preclinical evidence to rethink the dietary approach to CKD management. While Pi restriction is not a new concept, relatively simple real-life changes to diet, such as avoiding phosphate-based food preservatives and increasing vegetable protein intake while decreasing meat intake, could be more strongly stressed. This could potentially provide efficacy in delaying progression of CKD in individuals with modifiable high Pi intake. Clinical trials are needed to demonstrate this.

## Acknowledgement

We would like to thank Jyoti Arora from the Division of Biostatistics at Washington University School of Medicine for advice in statistical analysis. The content is solely the responsibility of the authors and does not necessarily represent the official views of the National Institutes of Health.

## Statement of Ethics

The Institutional Animal Care and Use Committee at Washington University in Saint Louis approved all animal studies (protocol ID 24-0117), and they adhere to the NIH Guide for the Care and Use of Laboratory Animals.

## Author Contributions

MJW and KAH designed the study. MJW, CBH, and HMP performed the experiments. MJW, HMP, and KAH conducted data analysis. MJW and KAH prepared, and all authors reviewed, the manuscript.

## Data Availability Statement

The data that support the findings of this study are openly available in Washington University School of Medicine’s Institutional Repository - Digital Commons Data@Becker, at http://doi.org/10.17632/mv3s66nkmb.1, reference number DOI: 10.17632/mv3s66nkmb.1.

